# High-Throughput Human Primary Cell-Based Airway Model for Evaluating Influenza, Coronavirus, or other Respiratory Viruses *in vitro*

**DOI:** 10.1101/2020.05.23.112797

**Authors:** A.L. Gard, R. Maloney, B.P. Cain, C.R. Miller, R.J. Luu, J.R. Coppeta, P. Liu, J.P. Wang, H. Azizgolshani, R.F. Fezzie, J.L. Balestrini, B.C. Isenberg, R.W. Finberg, J.T. Borenstein

## Abstract

Influenza and other respiratory viruses represent a significant threat to public health, national security, and the world economy, and can lead to the emergence of global pandemics such as the current COVID-19 crisis. One of the greatest barriers to the development of effective therapeutic agents to treat influenza, coronaviruses, and many other infections of the respiratory tract is the absence of a robust preclinical model. Preclinical studies currently rely on high-throughput, low-fidelity *in vitro* screening with cell lines and/or low-throughput animal models that often provide a poor correlation to human clinical responses. Here, we introduce a human primary airway epithelial cell-based model integrated into a high-throughput platform where tissues are cultured at an air-liquid interface (PREDICT96-ALI). We present results on the application of this platform to influenza and coronavirus infections, providing multiple readouts capable of evaluating viral infection kinetics and potentially the efficacy of therapeutic agents in an *in vitro* system. Several strains of influenza A virus are shown to successfully infect the human primary cell-based airway tissue cultured at an air-liquid interface (ALI), and as a proof-of-concept, the effect of the antiviral oseltamivir on one strain of Influenza A is evaluated. Human coronaviruses NL63 (HCoV-NL63) and SARS-CoV-2 enter host cells via ACE2 and utilize the protease TMPRSS2 for protein priming, and we confirm expression of both in our ALI model. We also demonstrate coronavirus infection in this system with HCoV-NL63, observing sufficient viral propagation over 96 hours post-infection to indicate successful infection of the primary cell-based model. This new capability has the potential to address a gap in the rapid assessment of therapeutic efficacy of various small molecules and antiviral agents against influenza and other respiratory viruses including coronaviruses.

## Introduction

The specter of a pandemic respiratory virus such as influenza or the current outbreak of coronavirus represents one of the greatest threats to human health [1,2]. For influenza, the development of efficacious, long-lasting, and seasonally-consistent vaccinations has been challenged by the occurrence of both antigenic drift and shift [3]. Currently, four antivirals are in wide use for treatment of influenza, three neuraminidase inhibitors and more recently, a cap-dependent endonuclease inhibitor, baloxivir marboxil [4]. These treatments are generally effective, but genetic changes to endemic viruses present challenges similar to those encountered during the development of vaccines [5]. These challenges are exacerbated by the long developmental timelines typically associated with manufacturing, scale up, and regulatory approval for novel antivirals or vaccinations. The current COVID-19 pandemic of the novel coronavirus has spurred aggressive vaccine development efforts, but even with this accelerated timeline, the current wave of infection is projected to yield significant morbidity and mortality [6]. Therefore, a critical element of medical treatment strategy is the application of repurposed FDA-approved antiviral medications originally developed for other diseases [7].

Several preclinical research tools are available to model respiratory viruses, but due to the complexity of the human lung and requirements to accurately model species-specific viral infections, these models provide a poor correlation as a basis for guiding clinical strategy [8]. For example, a standard approach to investigate influenza A viruses (IAV) typically begins with the inoculation of cell lines in conventional culture well plates, such as Madin-Darby canine kidney (MDCK) or Calu-3 cells [9,10]. While these cells infect easily and provide a rapid early assessment of the potential efficacy of a therapeutic for IAV, they lack the critical mechanisms and signaling pathways that healthy human airway tissues provide in response to viral infection. Therefore, although these systems are easy to use, they are not often accurate predictors of human clinical efficacy [11]. For coronaviruses, Calu-3 or Vero E6 cells are often used for similar reasons, but again differ by species, cell source, disease state, and innate antiviral signaling pathways [12,13] compared to normal, healthy human conducting airway and alveolar epithelium [14], thus providing an insufficient model. Unfortunately, complications also exist within animal models of IAV, not only in mice, rats, and guinea pigs, but even in more human-relevant options for evaluation such as ferrets and macaques [15,16]. These models often require species-adapted viral strains, and fail to recapitulate features of human innate antiviral responses and clinical symptoms of infection [17,18]. Similar challenges are faced for coronavirus infection studies, where the low-throughput and limited availability of appropriate animal models, combined with concerns about their predictive power, are forcing early and often high-risk testing in humans. While many experimental studies are underway to identify drugs with antiviral activity against SARS-CoV-2 [19], the discovery pipeline would benefit significantly from rapid and accurate preclinical models based on human cell culture.

The challenges with preclinical models and tools have spurred the development of cell culture platforms comprising human primary cells in a microfluidic environment, commonly known as microphysiological systems (MPS) or organs-on-chips [20]. Over the past two decades, these devices have evolved from early toxicology platforms to today’s wide range of technologies including multi-organ systems [21,22] and a variety of organ-specific models including airway or alveolar tissue systems [23,24]. While the integration of human primary cells or stem cell-derived populations in a physiologically-relevant microenvironment represents a tremendous advance over the use of cell lines in principle, in practice the application of these systems in pharmacological research and development has been severely gated by several factors. These limitations include low-throughput and complex operation, a lack of relevant metrics for system assessment, the use of research-grade materials and components, the lack of critical components of tissue or organ structure required to replicate infection, and most critically, an absence of confidence in the *in vitro-in vivo correlation* (IVIVC) for these technologies. Our group has developed systems that offer higher throughput, compatibility with existing pharmaceutical laboratory infrastructure, convenient readouts for quality control and physiological monitoring, and that are demonstrating the potential as platforms for preclinical studies across many disease areas and application domains [25,26].

In this manuscript, we build on our previous work and the work of others to develop an *in vitro* model with an ALI capability sufficient to support physiologically relevant epithelial differentiation such as that in gut or kidney models [25,26]. For functional proximal airway or alveolar models, human primary cells should be cultured at an ALI representing a suitable analogue for the respiratory barrier tissues that mimics mucus formation, mucociliary flow, and a range of physiologically relevant responses [23,24]. Our airway model, established in the PREDICT96-ALI platform, provides this ALI and supports IAV and coronavirus infection, while enabling high-throughput, real-time evaluation of experiments across various viral strains, multiplicities of infection (MOIs), human donors, time points, and statistically significant numbers of replicates on a single 96-device engineered platform. Three strains of IAV, A/WSN/33 H1N1, A/California/04/09 H1N1, and A/Hong Kong/8/68 H3N2 are used to inoculate human primary airway tissues cultured at an ALI, and viral infection kinetics are monitored through a combination of readouts including quantitation of viral load. As a proof-of-concept, viral infection kinetics are investigated for the A/WSN/33 H1N1 strain of IAV in response to dosing with oseltamivir, demonstrating how this system might be applied to evaluate the efficacy of antiviral therapeutic compounds. We also confirm expression of the receptor ACE2 and the serine protease TMPRSS2 in these tissues, and we demonstrate viral propagation and infection kinetics of HCoV-NL63, which, like SARS-CoV and SARS-CoV-2, utilizes ACE2 as its target receptor. These results demonstrate that the PREDICT96-ALI platform can be used to model IAV and coronavirus infections and could potentially be used to screen the efficacy of candidate therapeutics in a clinically relevant manner.

## Methods

### Microfluidic Platform

The PREDICT96 organ-on-chip platform consists of a microfluidic culture plate with 96 individual devices and a perfusion system driven by 192 microfluidic pumps integrated into the plate lid [25,26]. Here the platform is adapted to support culture of barrier tissues, such as airway tissues, by establishment and maintenance of an ALI.

Each device in the PREDICT96-ALI culture plate was formed from a 2×2 array within a standard 384-well plate, with the wall between the top two wells removed to create a 3-well cluster. This configuration is shown in **Figure 1a**. A slotted hard plastic microwell (0.864 mm height, 1 mm width, 2.5 mm length) was positioned in the center of the large well, with a microfluidic channel (0.25 mm height, 1 mm width) aligned underneath that was linked to the two remaining wells in each device. The channel was capped from below with a thin optically clear layer. A microporous membrane was used to separate the top microwell from the bottom microchannel to permit the establishment of an ALI.

**Figure 1.**
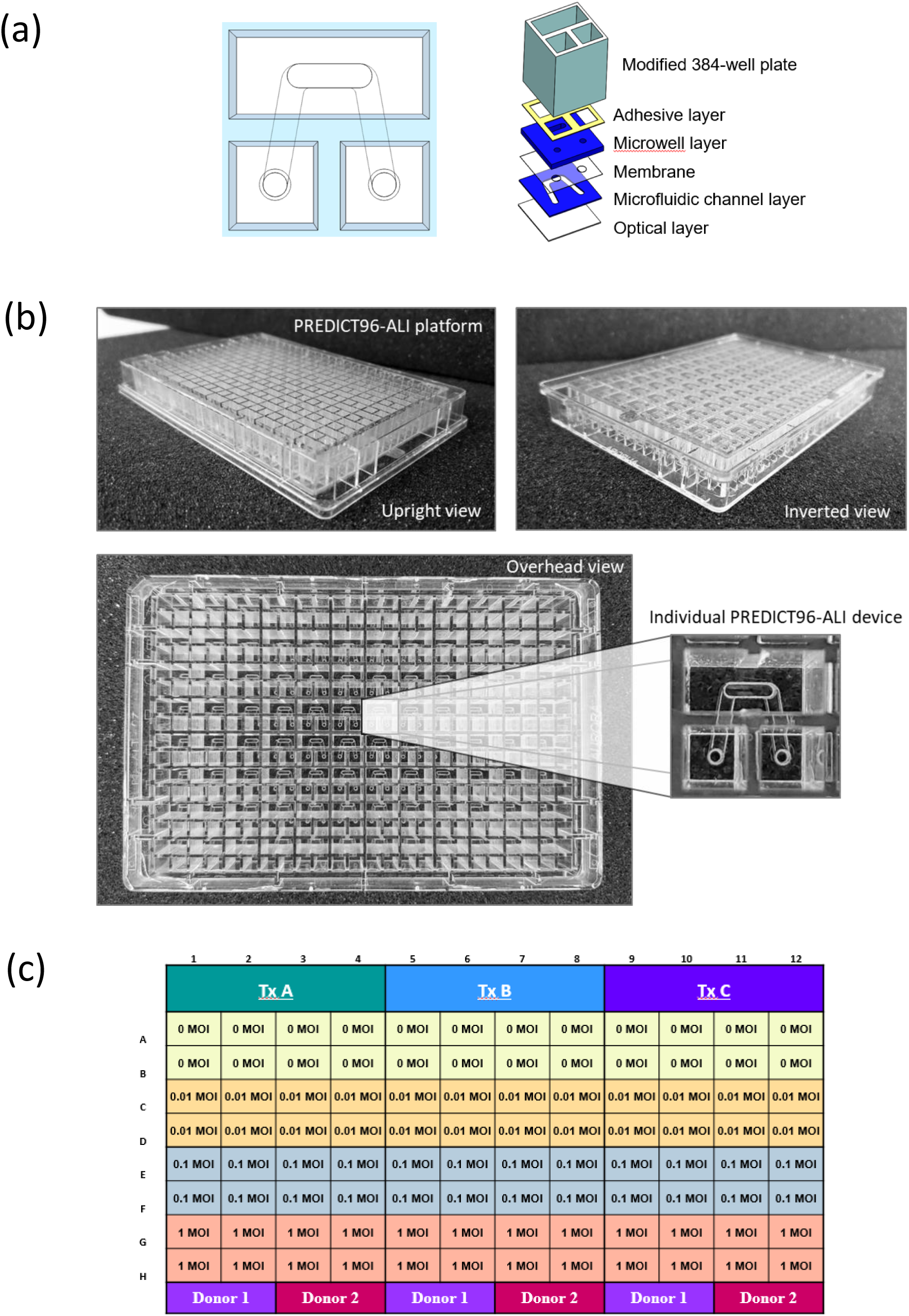
PREDICT96-ALI Platform. **(a)** Schematics illustrating the configuration of the PREDICT96-ALI platform, where the top chamber appears as an oval region and the underlying bottom chamber an inverse U-shaped channel with inlet and outlet ports, and the stack of polymer films used to create the structure for each of the 96 devices from a 2×2 array within a standard 384-well plate. **(b)** Photographs of the upright view (top left), inverted view (top right), and overhead view (bottom) of the PREDICT96-ALI platform configuration showing 8 rows and 12 columns of the individual device structure shown in panel **a. (c)** Sample plate map showing a potential experimental design where three therapeutics (A,B,C) are evaluated across two donor populations of NHBE cells (1,2), each at three MOIs (0.01, 0.1 and 1) and with four replicates per each condition.

To fabricate the PREDICT96-ALI culture plate, a UV laser system (Protolaser U4: LPKF Laser and Electronics, Garbsen, Germany) was used to laser-cut thin films of cyclic olefin polymer (COP) (ZF14-188: Zeon Corp., Tokyo, Japan) and cyclic olefin copolymer (8007 COC: Tekni-plex, Wayne, PA, USA). The COP layers were adhered together using low-glass transition temperature COC in a heated hydraulic press (Carver Inc., Wabash, IN, USA), and were separated with a 24 µm-thick track-etched polycarbonate membrane with pore diameter of 0.4 or 1 µm (it4ip S.A., Louvain-la-Neuve, Belgium) patterned with an array of holes to provide fluidic access ports to the bottom channel. The microfluidic stack is attached to a modified 384-well COP plate (Aurora Microplates, Whitefish, MT, USA) using a 0.135 mm thick pressure sensitive adhesive consisting of a 25.4 µm polyester carrier film with MA-69 acrylic medical grade adhesive ARcare® 90106 (Adhesives Research, Inc., Glen Rock, PA, USA), which had been previously laser-cut in the pattern of the well plate grid and laminated with a hydraulic press at 1.0 MPa for 2 min. Further details on the fabrication process are found elsewhere [25,26].

### Integrated Micropumps

As described in our previous work [25], in order to establish recirculating flow in the microchannels, we incorporate a self-priming micropump array into the lid that serves as the fluidic interface with the culture plate via stainless steel tubes. The PREDICT96 pumping system has 192 individual pneumatically-actuated micropumps embedded in the plate lid: two per culture device, one for the chamber above the membrane and one for below. However, since the upper chamber is at ALI, only the 96 pumps serving the microchannel are in use during experiments. Actuation of the pumps transfers media between the wells linked by the bottom channel and establishes a hydrostatic pressure differential, inducing flow through each microchannel.

The pneumatic and fluidic manifolds of the micropump array were constructed with laser-micromachined Ultem polyetherimide and Kapton polyimide films (McMaster-Carr, Elmhurst, Illinois USA) laminated in a heated hydraulic press with phenolic butyral thermosetting adhesive film (R/flex 1000, Rogers Corp., Chandler, Arizona, USA). Fluidic tubing (21G 316L stainless steel hypodermic tubes: New England Small Tube Corp., Litchfield, NH, USA) were glued into the assembly using 353NDPK Hi-Temp Epoxy (Thorlabs, Newton, NJ, USA).

### Culture, Cryopreservation, and PREDICT96-ALI Seeding and Differentiation of NHBEs

Normal Human Bronchial Epithelial Cells (NHBEs) were purchased from Lifeline Cell Technology and Lonza (**Table 1**). To maintain stocks of each NHBE donor, cells were thawed and plated at 2500 cells/cm^2^ on 804G media-coated tissue culture flasks (804G media made as described elsewhere [27]), cultured in Bronchialife (Lifeline Cell Technology), and passaged using Accutase (Sigma). After two passages, NHBEs were cryopreserved using 65% FBS (Thermo Fisher), 25% Bronchialife (Lifeline Cell Technology), and 10% DMSO (Sigma). Prior to seeding, PREDICT96-ALI plates were sterilized overnight with ethylene oxide gas followed by one week of outgassing in a vacuum chamber. PREDICT96-ALI plates were subsequently treated with plasma for 120 seconds and washed briefly with 70% ethanol, rinsed three times in distilled water, and coated overnight in 804G media. To seed PREDICT96-ALI plates, NHBEs were thawed, counted, re-suspended in complete small airway epithelial cell growth media (SAGM; Lonza), 100 U/mL penicillin-streptomycin (Thermo Fisher), 5 μM ROCKi (Tocris), 1 μM A-83-01 (Tocris), 0.2 μM DMH-1 (Tocris), 0.5 μM CHIR99021 (Tocris) as described elsewhere [28] (hereafter referred to as SAGM + 4i), and plated at 10,000 cells per device in a 3 μL volume directly onto the membrane. After 48 hours (h) of growth in SAGM + 4i media, differentiation was initiated using fresh HBTEC-ALI media (Lifeline Cell Technology) plus 100 U/mL penicillin/streptomycin. After 48 h of submerged differentiation, the ALI was initiated by aspirating media from the apical surface of the tissue in the top chamber, while 60 μL of fresh HBTEC-ALI or custom-ALI media was added to the bottom chamber. Pumping was initiated at 1 μL /min in the bottom channel, and the media in the bottom channel was changed daily thereafter. Tissues were matured over the course of 3-5 weeks in ALI culture prior to viral infection experiments. To remove accumulated mucus from the apical surface of the maturing tissue, 100 µL of 1x Hank’s Balanced Salt Solution (HBSS) was added to the apical surface of each tissue and incubated on the tissues for 1 h at 37 °C with rocking, subsequently followed by an additional 5 min wash with 100 µL of 1x HBSS at room temperature every 7 days (d). Mucus washings were pooled, collected and stored at −80°C until processed. Tissue maturity and quality control was scored by a combination of metrics including barrier function, percent ciliated cells, ciliary beat, mucus secretion, and global tissue morphology, and this score was used to determine if individual devices of PREDICT96-ALI airway tissue were suitable for downstream experimentation and viral infection. Downstream processing of tissues following viral inoculation was generally conducted at 4-6 weeks of ALI and included tissue apical washes, basal media collection, measurement of barrier function, harvest of tissue for RNA extraction, and fixation for immunofluorescence imaging.

**Table I.** Summary detailing the primary normal human tracheobronchial epithelial cells (NHBEs) used to grow human airway tissue within the PREDICT96-ALI platform. Vendor, catalogue number, batch number (donor lot), age, sex and race of each donor provided.

| Donor No. | Cell Type | Company | Catalogue No. | Batch No. | Age (yo) | Sex | Race |
| --- | --- | --- | --- | --- | --- | --- | --- |
| A | NHBE | Lonza | CC-2540S | 630564 | 16 | M | C |
| B | NHBE | Lifeline | FC-0035 | 05101 | 29 | F | C |

### Immunofluorescence and confocal imaging

To prepare PREDICT96-ALI airway tissues for immunofluorescence staining, tissues were fixed *in situ* using 4% paraformaldehyde (Sigma) for 15 min at room temperature rocking and washed three times with 1x phosphate buffered saline (PBS). Tissues were subsequently permeabilized with 0.3% Triton-100 (Sigma) for 30 min and blocked with 3% normal goat serum (NGS, Thermo Fisher) containing 0.1% Tween-20 (Sigma) for 60 min at room temperature. Primary antibodies for basal cells (CK5, Thermo Fisher), goblet cells (Muc5AC, Thermo), ciliated cells (acetylated-tubulin, Abcam and beta-tubulin, Sigma), cellular proliferation (Ki67, Abcam), and influenza A nucleoprotein (IAV-NP, Abcam) were diluted 1:100 in 3% NGS including 0.1% Tween-20 and incubated overnight at 4°C rocking. Devices were washed twice with PBS followed by one rinse with 3% NGS plus 0.1% Tween-20 for 5 min each with rocking and secondary antibodies Alexa Fluor 488 goat anti-mouse IgG (Thermo Fisher) and Alexa Fluor 555 goat anti-rabbit IgG (Thermo Fisher) were diluted 1:300 and incubated for 18 h at 4 °C with rocking. Secondary antibodies were removed by washing three times with PBS containing 0.1% Tween-20, and tissues were incubated with rocking at room temperature for 30 min with Hoechst 33342 nuclear stain (Thermo Fisher, 10 mg/mL stock) at a 1:800 dilution and Phalloidin-iFluor 647 Reagent (Abcam) at 1:1000. After 30 min, the PREDICT96-ALI tissues were washed at least three times for 5 min with PBS prior to imaging. Stained PREDICT96-ALI devices were imaged using a Zeiss LSM700 laser scanning confocal microscope and Zen Black software. Tile scans and z-stacks of the tissue at 20x and 40x magnification were acquired.

### Viral Stocks, Propagation, Titer and Anti-Viral Compound

Influenza A viral strains A/WSN/33 H1N1 (influenza A/WSN/33, WSN/33, gift of Dr. Peter Palese), A/California/04/09 H1N1 (ATCC), and A/Hong Kong/8/68 H3N2 (Charles River). IAV strains were propagated in MDCK cells (ATCC), harvested in Dulbecco’s Modified Eagle Medium (DMEM, Thermo Fisher) containing 7.5% bovine serum album (BSA, Invitrogen), 100 U/mL penicillin-streptomycin (Invitrogen), and 0.1 µg/mL N-tosyl-L-phenylalanine chloromethyl ketone-trypsin (TPCK-trypsin, Sigma) and titered by viral plaque assay using MDCK cells as previously reported [29]. Viral infection experiments using PREDICT96-ALI tissue used A/WSN/33 H1N1 at passage 1, A/California/04/09 H1N1 at passage 3, and A/Hong Kong/8/68 H3N2 at passage 2.

Human coronavirus strain (HCoV) NL63 was purchased from ATCC and propagated two times in Lilly Laboratories Cell-Monkey Kidney 2 (LLC-MK2) using a method adapted from Herzog [30]. In brief, HCoV-NL63 was grown using LLC-MK2 cells in a limiting dilution series. Recovered virus was harvested from the last well of the dilution series showing diffuse cytopathic effect (CPE) at 4-5 d post-infection (p.i.). Subsequently, confluent LLC-MK2 monolayers were inoculated in 25 cm^2^ flasks with a 1:10 dilution of viral supernatant from the prior viral passage. The flasks were incubated at 34°C, 5% CO_2_, and harvested on day 4 when diffuse CPE was observed. To harvest the virus, flasks were frozen at −80 °C and thawed at 37 °C twice, and cells and supernatant were pooled and centrifuged at 4 °C for 10 min at 3000 rpm. Cleared supernatant was aliquoted and stored at −80 °C, and is referred to henceforth as passage 2 virus. HCoV-NL63 was harvested in EMEM (ATCC) containing 0.1% BSA (Sigma) and 100 U/mL penicillin-streptomycin (Thermo Fisher). Viral infection experiments with PREDICT96-ALI tissue used HCoV-NL63 at passage 2.

Viral plaque assay using LLC-MK2 cells were used to titer propagated HCoV-NL63. A 10-fold serial dilution was performed using the propagated virus, followed by 1 h of viral binding (100 µL of appropriate viral dilution per well) with rocking at 34 °C on confluent LLC-MK2 cells. After unbound virus was removed with one wash of phosphate-buffered saline (PBS), the cells were overlaid with agar (0.3%) in EMEM supplemented with 0.1% BSA and 100 U/mL penicillin-streptomycin. After the agar solidified at room temperature, the plates were incubated for 6 d at 34 °C until diffuse CPE was observed. The gel overlay was removed and the wells washed once with PBS. Cells were fixed using 4% paraformaldehyde (Sigma) over night at 4°C and stained with 0.1% crystal violet (Sigma) in 4% ethanol (Sigma) for 30 min at room temperature. Stained cells were washed in PBS and imaged, and plaque counts were determined using images analyzed using a customized MATLAB script to determine the plaque forming units (PFU) per mL.

Oseltamivir carboxylate (F. Hoffmann-La Roche Ltd., Basel, Switzerland) was used in anti-viral screens performed on PREDICT96-ALI tissue. Oseltamivir was diluted to 1 µM in complete HBTEC-ALI or custom-ALI media and applied to the bottom channel of PREDICT-ALI airway tissues subject to anti-viral evaluation following the standard apical wash of the tissue and 2 h prior to IAV-inoculation. Upon and after IAV-inoculation of the apical side of the PREDICT96-ALI tissue, 1 µM oseltamivir was maintained in the basal media for the duration of the study. Basal media changes including oseltamivir occurred every 24 h.

### Infection of PREDICT96-ALI Tissues with Viral Strains

To prepare PREDICT96-ALI tissues for influenza and coronavirus infection, a mucus wash using 1x HBSS as described above was performed. Viral strains were thawed on ice and individually diluted in viral infection media to reach the relevant MOI. Influenza virus infection media was composed of a single IAV strain diluted to the appropriate MOI in HBSS for A/WSN/33 and in HBSS including 1 µg/mL TPCK-trypsin (Sigma) for A/California/04/09 and A/Hong Kong/8/68. Coronavirus infection media was composed HCoV-NL63 diluted to the appropriate MOI in HBSS. When the mucus wash was completed, viral inoculum was added to the apical side of the tissues. Each experiment included both an untreated control in which ALI was maintained and an untreated control that was submerged in HBTEC-ALI or custom-ALI media for comparison to the viral inoculum-treated groups. The time of incubation varied by viral strain, with A/WSN/33 and HCoV-NL63 each individually incubated on the PREDICT96-ALI tissues for 6 h with rocking and A/California/04/09 and A/Hong Kong/8/68 each individually incubated for 1 h with rocking. Following incubation, both the top chamber and the bottom channel were aspirated, the apical surface of the tissue washed three times with 1x HBSS with the final wash collected for reference, and this was followed by complete aspiration of the top chamber to resume ALI culture. A volume of 60 μL of HBTEC-ALI or custom-ALI media was added to the bottom channel to resume normal culture conditions, with 1 µg/mL TPCK-trypsin maintained in the media of devices that had been inoculated with A/California/04/09 and A/Hong Kong/8/68. At 24 or 48 h intervals p.i., an apical wash using 1x HBSS was performed at 34-37 °C for 1.5 h with rocking to collect both mucus and virus. Specifically, 100 µL of 1x HBSS was added to the apical surface of each tissue and incubated on the tissues for 45 min at 34-37 °C with rocking, subsequently followed by an additional 50 µL wash with 1x HBSS for 45 min at 34-37 °C with rocking and a final 50 µL wash with 1x HBSS for 5 min at room temperature. Apical washings were pooled, collected and stored at −80 °C until processed. Basal media was collected at time points p.i. that correspond to mucus and virus wash collections and stored at −80 °C. Following apical wash and basal media collections, 60 μL of HBTEC-ALI or custom-ALI media (including 1 µg/mL TPCK-trypsin for devices that had been inoculated with A/California/04/09 and A/Hong Kong/8/68) was added to the bottom channel and complete aspiration of the top chamber to resume ALI culture was performed to resume normal culture conditions.

### RNA extraction and qRT-PCR

Supernatant viral RNA was collected from the apical side of PREDICT96-ALI tissues and isolated using a QIAamp Viral RNA Mini Kit (Qiagen) following the manufacturer’s specifications. Supernatant volumes of 100 μL were brought to 140 μL using HBSS. In order to collect bulk tissue RNA from devices, RLT buffer (Qiagen) with 0.01% v/v 2-mercaptoethanol (Sigma) was added to both the top chamber and bottom channel to disrupt the differentiated tissue. Tissue RNA was extracted from PREDICT96-ALI devices using an RNeasy Micro Kit (Qiagen) per the manufacturer’s instructions. One-step quantitative reverse transcription polymerase chain reaction (qRT-PCR) was then performed on extracted RNA samples using a QuantiTect Probe RT-PCR kit (Qiagen) following the standard protocol for a QuantStudio 7 Flex RT-PCR system. Briefly, 7.8 μL of the extracted supernatant viral RNA was used in a 20 μL reaction volume and 3.8 μL tissue RNA was used in a 20 μL reaction volume, and samples were run in duplicate. The reaction was run in an Applied Biosystems QuantStudio 7 Flex System (Thermo Scientific) using the following condition: 50 °C for 20 min, 95 °C for 5 min, 40 cycles of 95 °C for 15 sec and 60 °C for 45 sec. Taqman primers and probe targeting IAV-M were ordered from Thermo Scientific with the following sequences: FLUAM-7-F: CTTCTAACCGAGGTCGAAACGTA, FLUAM-161-R: GGTGACAGGATTGGTCTTGTCTTTA, FLUAM-49-P6: TCAGGCCCCCTCAAAGCCGAG, and Taqman primers and probe targeting HCoV-NL63 (Assay ID: Vi06439673_s1) were ordered from Thermo Scientific. Taqman Gene Expression Assays (Thermo Scientific) were used to target the following proteins in lung tissue: ACE2 (Hs01085333_m1), TMPRSS2 (Hs01122322_m1), TUBB6 (Hs00603164_m1), and Muc5AC (Hs01365616_m1). Absolute quantification (copies/mL) of supernatant viral RNA was calculated using a standard curve generated from serial dilutions of A/PR/8/34 viral RNA (Charles River Laboratories) or linearized HCoV-NL63 viral RNA (ATCC). Comparative cycle threshold (Ct) values were determined using the method described by Schmittgen and Livak [31] using normalization to the housekeeping gene GAPDH.

### Statistical analysis

Data are presented as mean ± standard deviation and were analyzed using GraphPad Prism version 8.3.1 for Windows (GraphPad Software). Statistical significance was determined using two-way analysis of variance (ANOVA) with Tukey’s or Sidak’s post hoc test for multiple comparisons, where appropriate. A p-value lower than 0.05 was considered statistically significant and is indicated in figures as follows: * P ≤ 0.05, ** P ≤ 0.01, *** P ≤ 0.001, and **** P ≤ 0.0001.

## Ethics statement

No ethics approval was required for this work.

## Results

### Creation of a high-throughput ALI platform for dynamic culture of human airway tissue: PREDICT96-ALI

The adaptation of the PREDICT96 platform [26] for the ALI model allows for the ability to inoculate cultured tissues in each channel apically or basally, to sample the basal media at regular intervals, and to examine the tissues at the completion of each study, as shown in **Figure 1a**. The schematic (left) shows the individual device configuration within the 96-device platform, where a 2×2 array from the 384 well standard structure is used to access the top chamber (top two wells of the 2×2 array) and the bottom channel is accessed by an inlet and outlet from the bottom two elements of the 2×2 array, respectively. An exploded-view schematic (right) of a single device configuration depicts the assembly of the material layers required to establish the microfluidic structure of the PREDICT96-ALI platform. Photographs in **Figure 1b** depict the PREDICT96-ALI platform from an upright view (top left), inverted view (top right), and overhead view (bottom), with 8 rows and 12 columns of the configuration shown in **Figure 1a. Figure 1c** presents a typical plate layout, highlighting the capacity of the 96-device plate to house replicates (four per condition) for each of three therapeutics, across two donor NHBE cell populations at three MOIs. The PREDICT96-ALI layouts can easily accommodate different experimental designs, for example replacing three therapeutic candidates with a test of three different human cell donors on the plate for a single therapeutic under evaluation.

### PREDICT96-ALI recapitulates healthy human airway structure and composition

First, the PREDICT96-ALI system was used to establish the healthy baseline airway model, prepared as described in the Methods section, and morphology and cell populations were evaluated using high resolution confocal microscopy and immunohistochemistry. An illustration depicts the development of a healthy airway model within PREDICT96-ALI (**Figure 2a**). The experimental timeline and setup for the system is outlined in **Figure 2b**. Fluorescent staining of ciliated cells, goblet cells, and basal cells (**Figure 2c**), as well as a cross-section of the pseudostratified epithelial layer (**Figure 2d**) are shown for a typical airway tissue within a single device of PREDICT96-ALI. Our observations for airway tissues in the PREDICT96-ALI platform regarding the relative populations of ciliated cells, basal cells, and goblet cells are consistent with our previously reported observations in earlier generations of our airway models [23].

**Figure 2.**
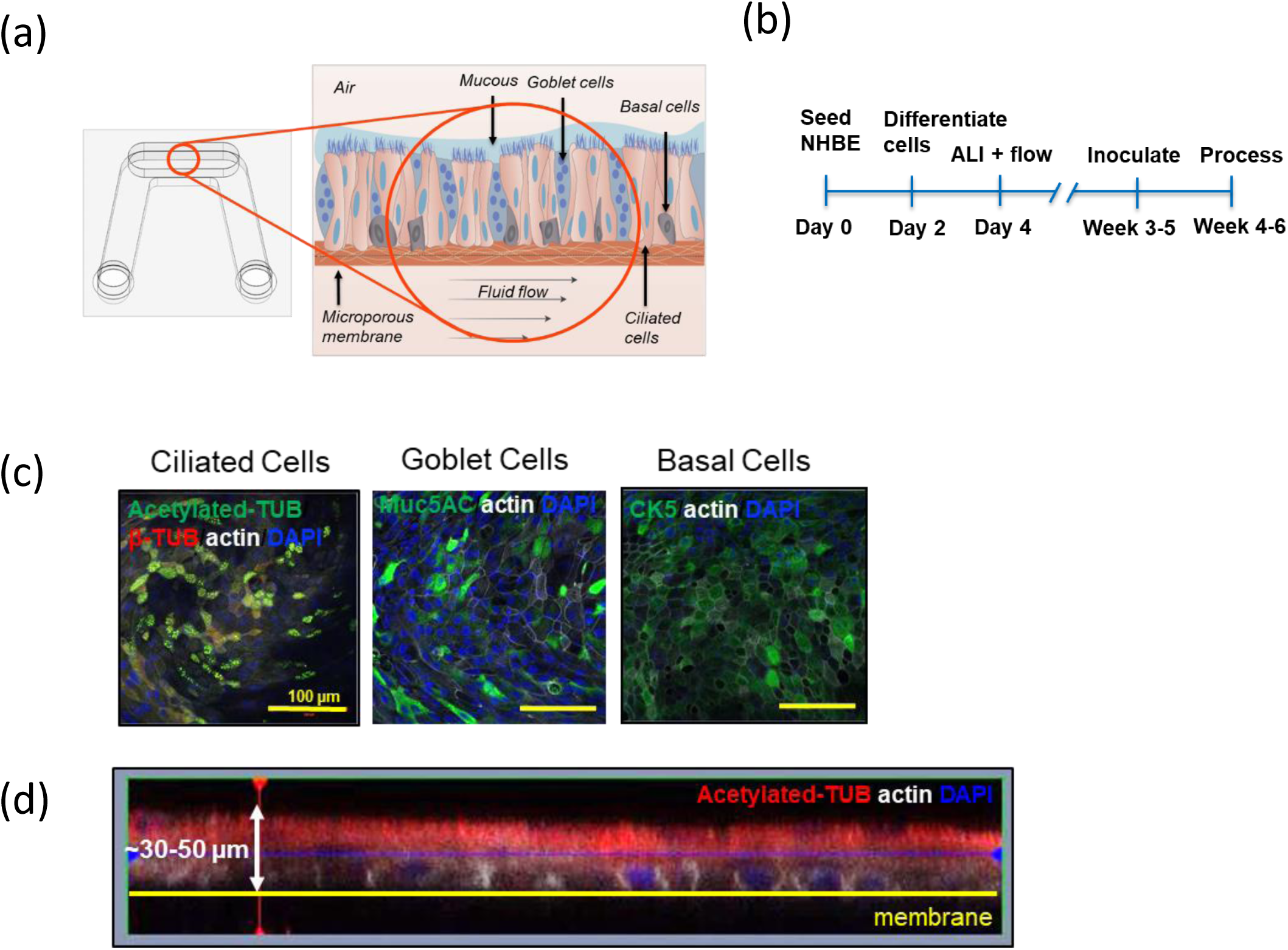
Healthy PREDICT96-ALI Airway Model. **(a)** Schematic illustrating the configuration of the biomimetic airway model within the PREDICT96-ALI platform with associated microenvironmental features including differentiated cell populations (ciliated, goblet, basal and club cells) of the mature tissue, apical mucus and periciliary fluid present on mature human airway, and basolateral fluid flow to recirculate nutrients, remove waste products, and oxygenate the media. **(b)** Timeline detailing proliferation, differentiation, inoculation and processing (sample collection, fixation, imaging, etc.) of the PREDICT96-ALI airway tissue over 4-6 weeks. **(c)** Staining of ciliated (acetylated-tubulin), goblet (Muc5ac) and basal (CK5) cells at 40x magnification shown, with a 100 μm scale bar. Donor B is featured as representative. **(d)** High-resolution confocal microscopy 40x z-stack orthogonal image of the pseudostratified epithelium with approximately 4 cell layers and 30-50 µm thick established after culture at 28 d at an ALI, with the membrane location shown for reference. Donor B is featured as representative.

### PREDICT96-ALI airway model supports infection with influenza A virus

Next, we examined the effect of inoculation of airway ALI cultures with various strains of IAV, including A/WSN/33 H1N1, A/California/04/09 H1N1, and A/Hong Kong/8/68 H3N2. In **Figure 3a**, immunofluorescence (IF) staining of ALI culture devices inoculated with various MOIs of A/WSN/33 are shown, demonstrating greater abundance of IAV nucleoprotein (NP) as the MOI increases. In **Figure 3b**, 20x images at the apical plane show IAV NP staining across the range of 0.1 – 10 MOI, with levels increasing substantially as the MOI rises from 0.1 to 1 and again from 1 to 10.

**Figure 3.**
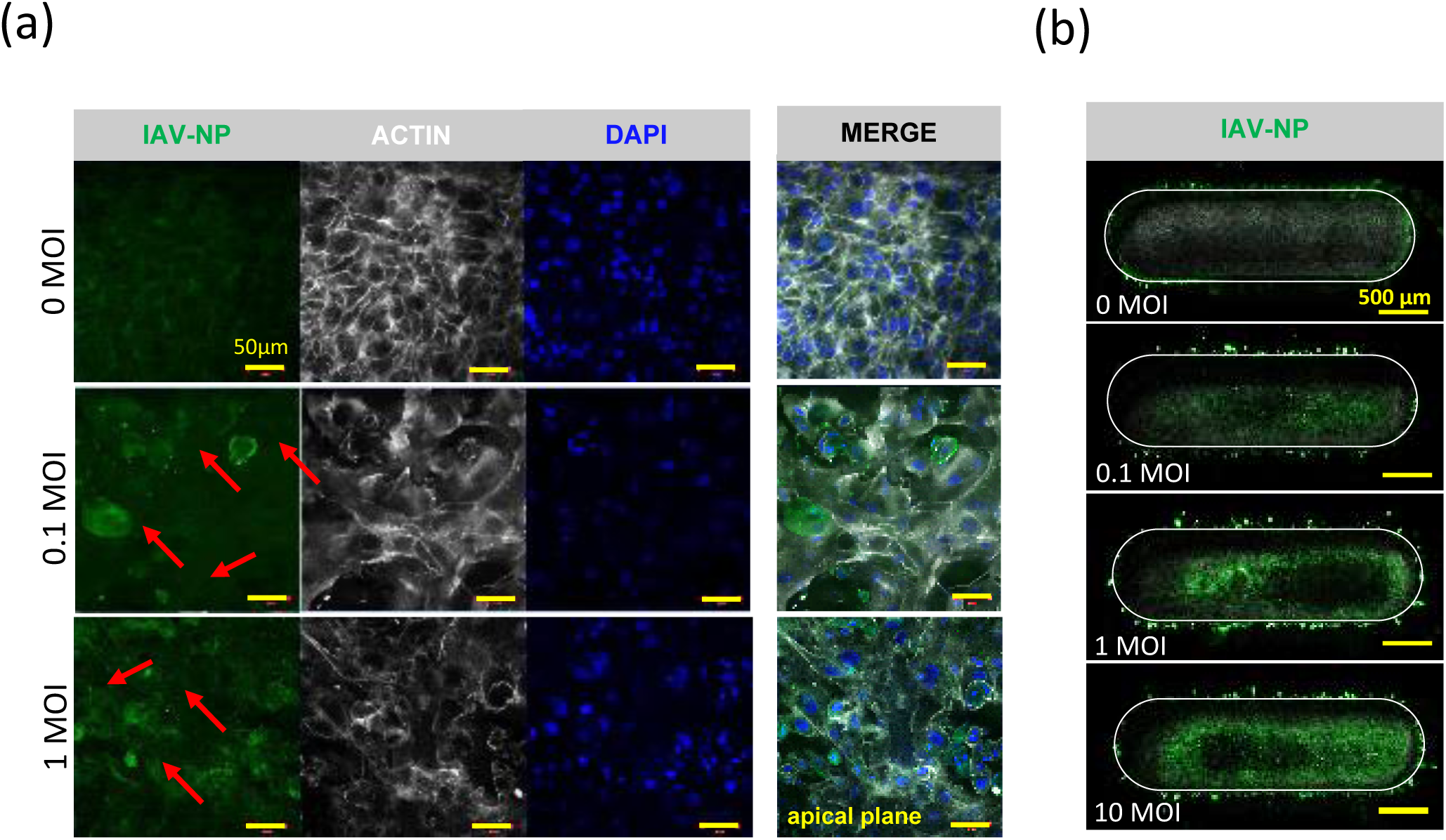
Immunofluorescence staining of IAV-infected PREDICT96-ALI Airway Tissue. **(a)** Staining of the nucleoprotein (NP, green) for IAV, actin (phalloidin, grey), and nucleic acids (DAPI, blue), including merged panels for all three stains, within PREDICT96-ALI airway tissue at 48 h p.i. following inoculation with A/WSN/33 (MOI 0, 0.1 or 1). Red arrows denote IAV-NP positive staining at MOI 0.1 and 1 as characterized in the apical plane of 40x images of the tissue. An increase in cytopathic effect (CPE) is observed with increasing MOI at 48 h p.i. Donor A is featured as representative. **(b)** Staining of the nucleoprotein (NP, green) captured as full device tile images of the apical plane of the tissue at 48 h p.i. and 20x magnification. Donor B is featured as representative.

Measurements of supernatant viral RNA were obtained using qRT-PCR, and are provided in the three panels in **Figure 4**. Viral copy numbers after inoculation at MOI of 0.1 and 1 were measured at 24 and 48 h p.i. for the A/WSN/33 strain of IAV (**Figure 4a**). In **Figure 4b** and **4c**, companion results for A/California/04/09 H1N1 and A/Hong Kong/8/68 H3N2 are shown, respectively. For the H1N1 and H3N2 strains, viral copies per mL rise monotonically with MOI, and continue to increase between 24 and 48 h p.i. For the WSN strain, viral copies per mL show a large increase at MOI = 1, and are relatively flat after 24 h p.i.

**Figure 4.**
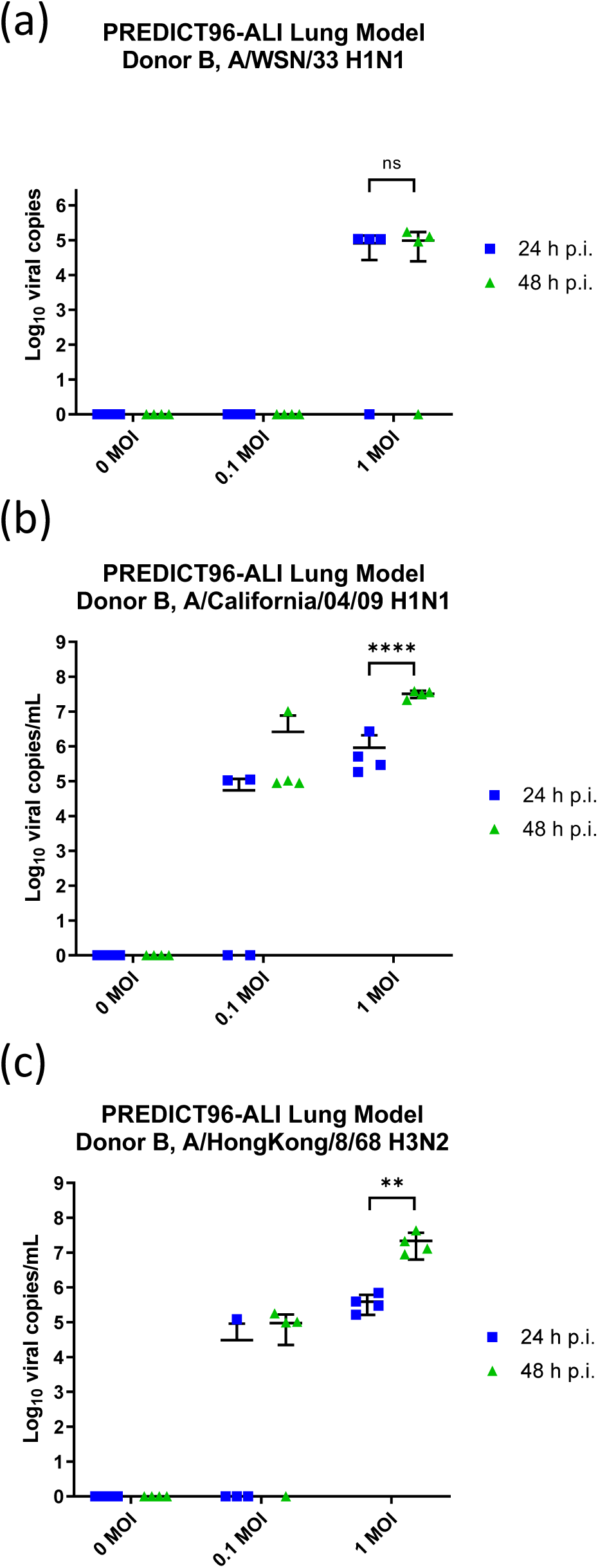
Infection Kinetics of IAV-inoculated PREDICT96-ALI Airway Tissue. qRT-PCR analyses for viral titer of the apical wash of PREDICT96-ALI airway tissue at 24 and 48 h (p.i.) **(a)** PREDICT96-ALI airway tissue inoculated with A/California/04/09 H1N1 and monitored for an increase in viral load at 24 h intervals with a statistically significant increase in viral titer at 48 h p.i. compared to 24 h p.i. for the MOI = 1 condition. **(b)** PREDICT96-ALI airway tissue inoculated with A/Hong Kong/8/68 H3N2 and monitored for an increase in viral load at 24 h intervals with a statistically significant increase in viral titer at 48 h p.i. compared to 24 h p.i. for the MOI = 1 condition. **(c)** PREDICT96-ALI airway tissue inoculated with A/WSN/33 and monitored for an increase in viral load at 24 h intervals, with non-significant (ns) changes in viral load observed for the MOI = 1 condition. Donor B is featured. Statistical significance: ** p ≤ 0.01 and **** p ≤ 0.0001. N = 2 independent experiments with data presented from one representative experiment. Per experiment, N = 3-4 tissue replicates per donor, time-point and condition

### Influenza A virus replication is reduced by oseltamivir in PREDICT96-ALI airway model

To determine if the PREDICT96-ALI airway model can be used to evaluate the efficacy of potential antiviral therapeutics, we investigated the effect of the antiviral agent oseltamivir (Tamiflu) – the most commonly used clinical anti-influenza therapy – for its ability to reduce viral load in IAV-inoculated PREDICT96-ALI airway tissue. The active form of oseltamivir, oseltamivir carboxylate, was introduced 2 h before viral inoculation and maintained in the bottom channel of the PREDICT96-ALI airway tissues during the course of infection with A/WSN/33 H1N1 virus at various MOIs. Oseltamivir (1 μM) significantly reduced influenza replication up to 48 h p.i. **(Figure 5)** relative to control tissue devices for the A/WSN/33 H1N1 strain at MOI = 1 and prevented virus-induced disruption to barrier function and epithelial tight junction formation (data not shown). Measurements for viral copies made in the PREDICT96-ALI airway model reflect those made in clinical settings among patients exposed to influenza and treated with oseltamivir [32,33,34,35], suggesting that the PREDICT96-ALI airway model can serve as a preclinical tool to evaluate potential therapies for combating respiratory infections of the human airway.

**Figure 5.**
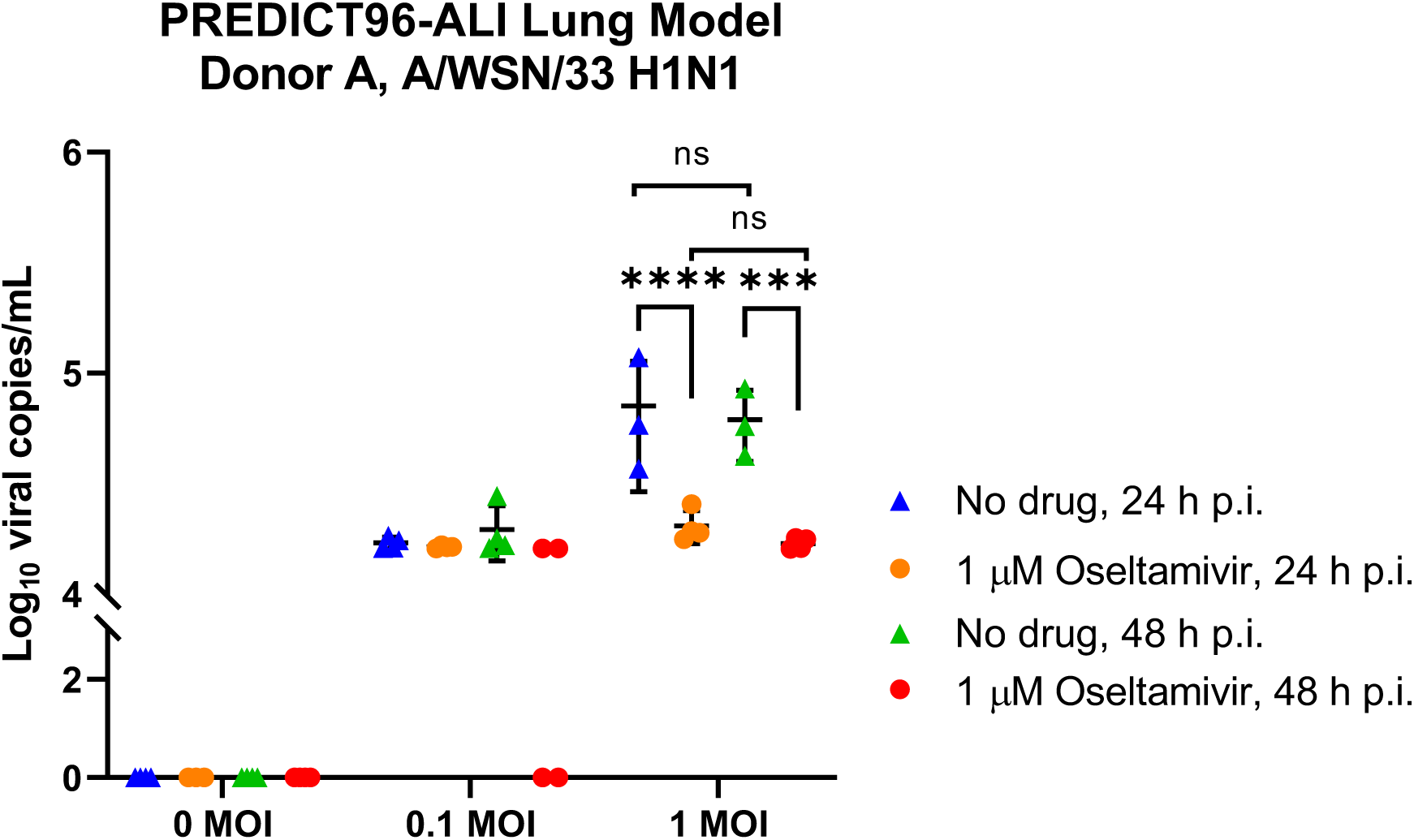
Infection Kinetics of IAV-inoculated PREDICT96-ALI Airway Tissue in Response to Oseltamivir. qRT-PCR analyses for viral copies of the apical wash of PREDICT96-ALI airway tissue at 48 h p.i. in absence (triangle, blue and green) or presence (circle, orange and red) of oseltamivir (Tamiflu) dosing at 1 μM at 24 (blue and orange) and 48 h p.i. (green and red). PREDICT96-ALI airway tissue inoculated with A/WSN/33 H1N1 at MOI 0, 0.1 and 1, and monitored for viral load over 24 h intervals up to 48 h p.i. with a statistically significant decrease in viral titer at MOI = 1 in response to oseltamivir dosing (circle) versus no drug (triangles) at both 24 and 48 h p.i.. Statistical significance: *** p ≤ 0.001 and **** p ≤ 0.0001. N = 2 independent experiments with data presented from one representative experiment. Per experiment, N = 3-4 tissue replicates per donor, time-point and condition

### ACE2 and TMPRSS2 transcript expression in PREDICT96-ALI airway model

In preparation for testing the viral respiratory infection model with coronaviruses, we probed for transcript expression of cell surface proteins ACE2 and TMPRSS2 known to play central roles in facilitating viral infection by SARS-CoV-2 [36]. Both HCoV-NL63 and SARS-CoV-2 gain entry into host cells via the ACE2 receptor, while TMPRSS2 facilitates SARS-CoV infection via cleavage mechanisms, and thus its presence is an important aspect of the model. Data for ACE2 and TMPRSS2 transcripts for cell populations derived from two different donors are plotted on a semi-logarithmic scale in **Figure 6**. Data from each donor evaluated suggest a lack of statistical difference in transcript expression in ACE2 and TMPRSS2 between donors. The consistency between these key cellular factors among different donors is encouraging, as it suggests that the platform is capable of supporting proliferation and maturation of tissues derived from numerous donors. Thus, evaluation of many donors can be accommodated in the PREDICT96-ALI system both as a healthy model and in the presence of pathogenic challenge.

**Figure 6.**
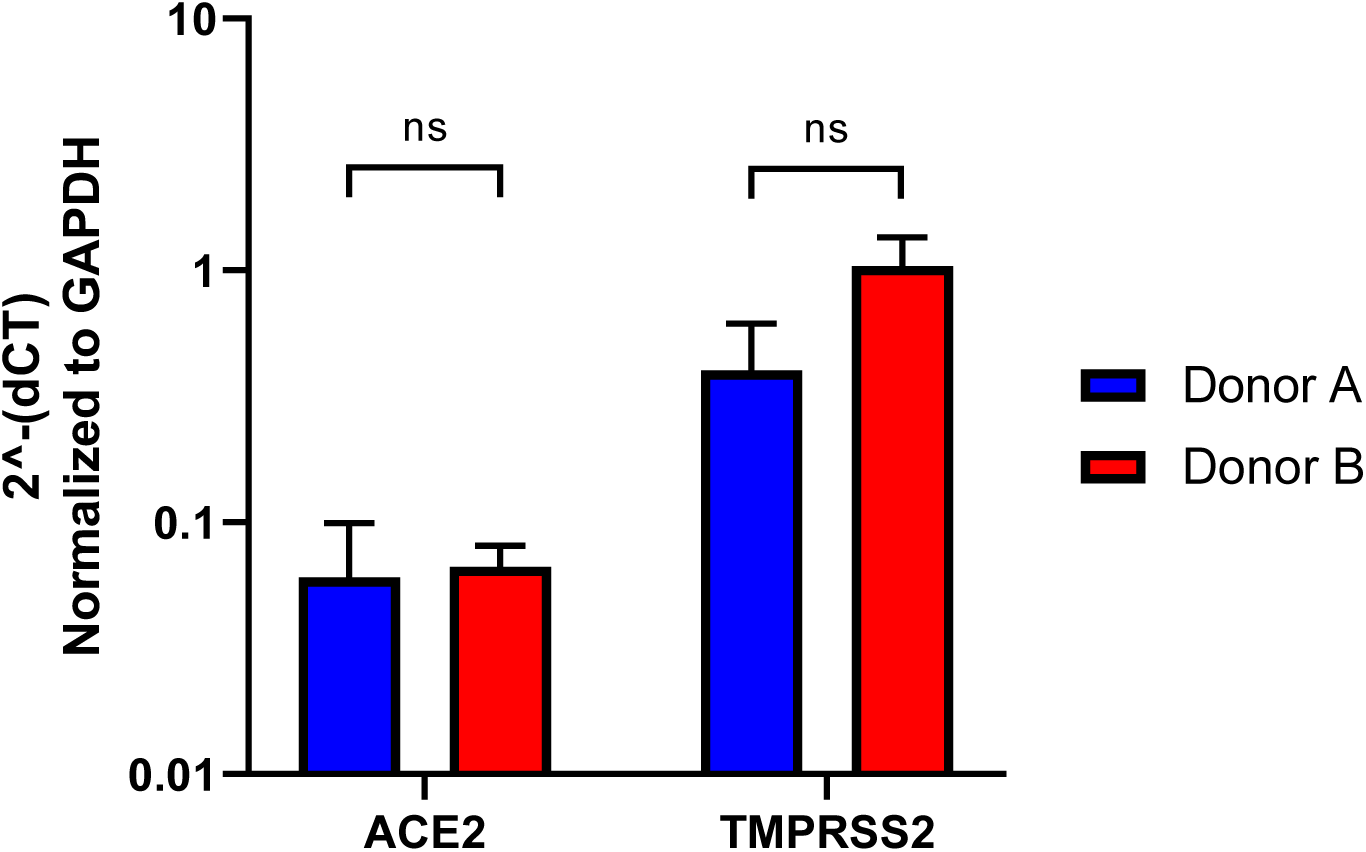
Comparative Quantification of ACE2 and TMPRSS2 in PREDICT96-ALI Airway Tissues. ACE2 and TMPRSS2, both critical for establishing HCoV-NL63 and SARS-CoV-2 infection, were detected by qRT-PCR from PREDICT96-ALI airway tissue. PREDICT96-ALI airway tissue from Donor A and B were matured to 4 weeks ALI and shown to have a non-significant (ns) difference in ACE2 and TMPRSS2 gene transcript levels when compared to one another. Comparative CT values were determined using the method described by Schmittgen and Livak [31] and used to determine the relative quantification of gene expression using GAPDH as a reference gene. N = 3-4 tissue replicates per donor.

### PREDICT96-ALI airway model supports infection with human coronavirus NL63

We have also investigated viral infection with HCoV-NL63 in the PREDICT96-ALI culture system with primary NHBEs, as shown in **Figures 7a** and **7b** for two different donor cell populations. In each case, viral copies in the supernatant were measured with qRT-PCR over periods extending to 96 h p.i. and across a range of MOIs. We observed a clear increase in viral load over these time periods for both donors, indicating the presence of viral propagation in the airway tissues. Viral copies rose sharply from 0 to 48 h for each MOI tested, while from 48 to 96 h the rate of viral propagation decreased and differed between the two donors.

**Figure 7.**
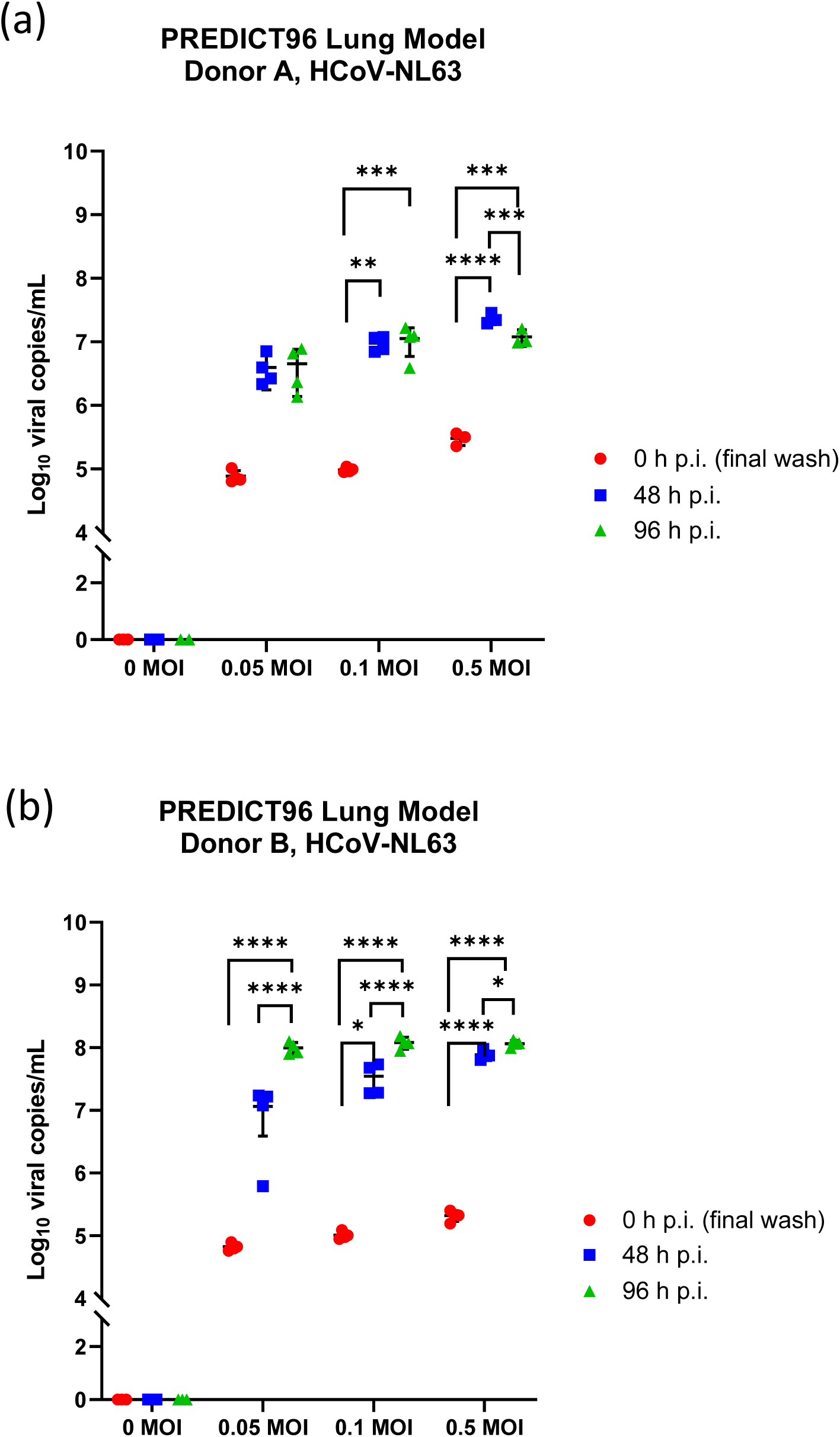
Infection Kinetics of HCoV-NL63-inoculated PREDICT96-ALI Airway Tissue. qRT-PCR analyses for viral titer of the apical wash of PREDICT96-ALI airway tissue at 0, 48 and 96 h p.i. for two human donors showing unique infection kinetic profiles. **(a)** PREDICT96-ALI Donor A airway tissue inoculated with HCoV-NL63 and monitored for an increase in viral load at 48 h intervals with a statistically significant increase in viral titer at 48 h p.i compared to 0 h p.i. for MOI 0.1 and 0.5, as well as a statistically relevant increase in viral titer at 96 h p.i. compared to 0 and 48 h p.i. for the MOI = 0.5 condition. ** p ≤ 0.01, *** p ≤ 0.001, **** p ≤ 0.0001. **(b)** PREDICT96-ALI Donor B airway tissue inoculated with HCoV-NL63 and monitored for an increase in viral load at 48 h intervals with a statistically significant increase in viral titer at 48 h p.i compared to 0 h p.i. for MOI 0.05, 0.1, and 0.5, as well as a statistically significant increase in viral titer at 96 h p.i. compared to 48 h p.i. for MOI 0.1 and 0.5 and compared to 0 h p.i. for the 0.05 MOI condition. Statistical significance: * p ≤ 0.05 and **** p ≤ 0.0001. N = 2 independent experiments with data presented from one representative experiment. Per experiment, N = 3-4 tissue replicates per donor, time-point and condition Different Y-axes reflect large difference in viral copy number detected in individual donors.

## Discussion

In this work, we leverage the PREDICT96-ALI platform to create a high-throughput airway infection model to assist in the development of therapeutics to treat infections. To do so, we developed a barrier tissue model at an ALI with human primary tracheobronchial cells, establishing a pseudostratified epithelium, and cultured atop a semipermeable membrane with independent flow control for each device in the bottom channel. The data presented demonstrate a robust model for the human airway using primary tracheobronchial epithelial cells, with an appropriate distribution of cell populations of ciliated, basal, and goblet cells and a pseudostratified epithelial layer morphology. This airway model was applied to the investigation of respiratory viral infections including IAV and HCoV-NL63, and infection kinetics were evaluated across several viral strains, donor populations, MOIs and time points. Experiments conducted with multiple donor populations of NHBE cells suggest the potential of this system for conducting studies of donor-dependent mechanisms of infection and responses to therapeutic interventions. Key advantages of this platform over existing *in vitro* platforms include the use of human primary tracheobronchial epithelial cells cultured at an ALI, a more physiologically relevant cell-to-media ratio than standard Transwell culture, integrated pumping of basal media to maintain precision control over solute concentration and to avoid the development of non-physiological gradients, and compatibility with high resolution and real-time imaging during dynamic pumping. While current organ-on-chip models possess some of these advantages, they are typically low-throughput systems that are designed for research environments and are often incompatible with workflows and standard instrumentation in pharmaceutical laboratories. Considering the throughput limitations of the competing organ-on-chip models, the PREDICT96-ALI platform offers an unmatched throughput that enables large combinations of conditions, such as evaluation of three therapeutics across four MOIs and time points with two viral strains and four replicates for each condition simultaneously on a single plate. The PREDICT96-ALI platform combines the high throughput, precision control and physiological relevance necessary to serve as a powerful tool for applications in disease modeling, drug development, and screening.

A critically important need in the field of respiratory virus research is the establishment of robust *in vitro* disease models that provide the precision, validation, throughput, and utility necessary for routine operation in drug development laboratories. Until recently, *in vitro* models suitable for the study of respiratory viral infections have been either high-throughput but simplistic 2D culture plate-based systems utilizing cell lines, or low-throughput and complex organ-on-chip models that are difficult to accommodate in the workflow of many laboratories. The PREDICT96 platform has been reported previously, and represents the first high-throughput organ-on-a-chip platform with integrated precision flow control in a standard ANSI SLAS (American National Standards Institute / Society for Laboratory Automation and Screening) well plate format [25]. Here, we deployed the airway tissue model successfully in the PREDICT96-ALI platform to investigate infections of three different strains of IAV; the A/WSN/33 strain and pandemic strains of H1N1 and H3N2. Infections were monitored using a combination of IF and qRT-PCR, across a range of MOIs and time points, providing a strong foundation for evaluation of therapeutic interventions. As a proof of concept, oseltamivir dosing of the PREDICT96-ALI airway model was shown to reduce viral copies of A/WSN/33 H1N1, demonstrating the potential of this system for screening therapeutic compounds for respiratory viruses.

Our human airway tissue model has applicability to respiratory viruses including coronaviruses, and we have demonstrated the relevance of this model to monitoring pathogen infection kinetics. To investigate the applicability of our model to novel viral pathogens, we examined the ability of HCoV-NL63, an alpha coronavirus sharing the same host entry receptor as SARS-CoV-2, to infect and replicate within PREDICT96-ALI.We both confirmed the presence of ACE2 and TMPRSS2 transcripts in the primary human cells within our model and successfully infected the PREDICT96-ALI airway model with HCoV-NL63 across a range of MOIs and for two different donor human primary epithelial cell populations. This demonstration of infection with HCoV-NL63 represents an important new capability for modeling coronavirus infections and potentially evaluating therapeutics, extending beyond current approaches that use cell lines in high-throughput systems or human primary lung cells in low-throughput Transwell formats. These data demonstrate the promise of the PREDICT96-ALI airway model as a new capability for preclinical evaluation of therapeutics for respiratory infections including SARS-CoV-2 in an efficient, robust, and high-throughput manner.

## Author Interests

The authors are all employees of Draper or the University of Massachusetts Medical School and declare no conflicts.

## Author Contributions

Experimental design was done by ALG, JW, RWF and JTB. Execution of the experiments was led by ALG along with RSM, CRM, RJL and PL. The PREDICT96 platform technology was adapted for ALI culture by BC, JRC, HA and BCI. The manuscript was written by ALG, RSM, BC, CRM, RJL, JLB, BCI and JTB, and reviewed and edited by all authors.

## Acknowledgements

The authors gratefully acknowledge Roger Odegard, Alla Gimbel, Melanie Trombly and David Paquette for their technical and programmatic support for this work, and Elizabeth Wiellette, Tim Petrie and Else Vedula for key technical guidance and critical review of the manuscript. Funding from two sources, Internal Draper Independent Research and Development (IR&D) for the coronavirus research, and from the DARPA BTO PREPARE program for the influenza A virus studies, is gratefully acknowledged. The work was partially supported by the PReemptive Expression of Protective Alleles and Response Elements (PREPARE) program from the Defense Advanced Research Projects Agency (DARPA), contracted via the Department of Navy, Office of Naval Research (ONR) under Federal Award Number N669911924036. The United States Government has a royalty-free license throughout the world in all copyrightable material contained herein. The views, opinions, and/or findings expressed are those of the author(s) and should not be interpreted as representing the official views or policies of the Department of Defense or the U.S. Government. Approved for Public Release. Distribution Unlimited.

## References

1 Taubenberger JK, Morens DM. 1918 Influenza: the mother of all pandemics. Emerg Infect Dis. 2006 Jan;12(1):15–22. doi: 10.3201/eid1201.050979. PMID: 16494711; PMCID: PMC3291398.

2 Paules CI, Marston HD, Fauci AS. Coronavirus Infections—More Than Just the Common Cold. JAMA. 2020;323(8):707–708. doi:10.1001/jama.2020.0757

3 Hyunsuh Kim, Robert G. Webster, and Richard J. Webby.Viral Immunology. Mar 2018.174–183.

4 Hayden FG, Sugaya N, Hirotsu N, et al. Baloxavir Marboxil for Uncomplicated Influenza in Adults and Adolescents. N Engl J Med. 2018;379(10):913–923. doi:10.1056/NEJMoa1716197

5 O’Hanlon R, Shaw ML. Baloxavir marboxil: the new influenza drug on the market. Curr Opin Virol. 2019 Apr;35:14–18. doi: 10.1016/j.coviro.2019.01.006. Epub 2019 Mar 8.

6 Lurie N, Saville M, Hatchett R, Halton J. Developing Covid-19 Vaccines at Pandemic Speed [published online ahead of print, 2020 Mar 30]. N Engl J Med. 2020;10.1056/NEJMp2005630. doi:10.1056/NEJMp2005630

7 Grein J, Ohmagari N, Shin D, et al. Compassionate Use of Remdesivir for Patients with Severe Covid-19 [published online ahead of print, 2020 Apr 10]. N Engl J Med. 2020;10.1056/NEJMoa2007016. doi:10.1056/NEJMoa2007016

8 Seok J, Warren HS, Cuenca AG, et al. Genomic responses in mouse models poorly mimic human inflammatory diseases. Proc Natl Acad Sci U S A. 2013;110(9):3507–3512. doi:10.1073/pnas.1222878110

9 Ilyushina, N. A., Ikizler, M. R., Kawaoka, Y., Rudenko, L. G., Treanor, J. J., Subbarao, K., & Wright, P. F. (2012). Comparative study of influenza virus replication in MDCK cells and in primary cells derived from adenoids and airway epithelium. Journal of virology, 86(21), 11725–11734. https://doi.org/10.1128/JVI.01477-12

10 Zeng, H., Goldsmith, C., Thawatsupha, P., Chittaganpitch, M., Waicharoen, S., Zaki, S., Tumpey, T. M., & Katz, J. M. (2007). Highly pathogenic avian influenza H5N1 viruses elicit an attenuated type i interferon response in polarized human bronchial epithelial cells. Journal of virology, 81(22), 12439–12449. https://doi.org/10.1128/JVI.01134-07

11 Takada, K., Kawakami, C., Fan, S. et al. A humanized MDCK cell line for the efficient isolation and propagation of human influenza viruses. Nat Microbiol 4, 1268–1273 (2019). https://doi.org/10.1038/s41564-019-0433-6

12 Desmyter J, Melnick JL, Rawls WE. Defectiveness of interferon production and of rubella virus interference in a line of African green monkey kidney cells (Vero). J Virol. 1968;2(10):955–961.

13 Mosca JD, Pitha PM. Transcriptional and posttranscriptional regulation of exogenous human beta interferon gene in simian cells defective in interferon synthesis. Mol Cell Biol. 1986;6(6):2279–2283. doi:10.1128/mcb.6.6.2279

14 Coleman CM, Frieman MB. Growth and Quantification of MERS-CoV Infection. Curr Protoc Microbiol. 2015;37:15E.2.1–15E.2.9. Published 2015 May 1. doi:10.1002/9780471729259.mc15e02s37

15 Groves, H. T., McDonald, J. U., Langat, P., Kinnear, E., Kellam, P., McCauley, J., Ellis, J., Thompson, C., Elderfield, R., Parker, L., Barclay, W., & Tregoning, J. S. (2018). Mouse Models of Influenza Infection with Circulating Strains to Test Seasonal Vaccine Efficacy. Frontiers in immunology, 9, 126. https://doi.org/10.3389/fimmu.2018.00126

16 Bouvier, N. M., & Lowen, A. C. (2010). Animal Models for Influenza Virus Pathogenesis and Transmission. Viruses, 2(8), 1530–1563. https://doi.org/10.3390/v20801530

17 Thangavel, R. R., & Bouvier, N. M. (2014). Animal models for influenza virus pathogenesis, transmission, and immunology. Journal of immunological methods, 410, 60–79. https://doi.org/10.1016/j.jim.2014.03.023

18 Seok J, Warren HS, Cuenca AG, et al. Genomic responses in mouse models poorly mimic human inflammatory diseases. Proc Natl Acad Sci U S A. 2013;110(9):3507–3512. doi:10.1073/pnas.1222878110

19 Costanzo M, De Giglio MAR, Roviello GN. SARS CoV-2: Recent Reports on Antiviral Therapies Based on Lopinavir/Ritonavir, Darunavir/Umifenovir, Hydroxychloroquine, Remdesivir, Favipiravir and Other Drugs for the Treatment of the New Coronavirus [published online ahead of print, 2020 Apr 16]. Curr Med Chem. 2020;10.2174/0929867327666200416131117. doi:10.2174/0929867327666200416131117

20 Low LA, Tagle DA. Microphysiological Systems (“Organs-on-Chips”) for Drug Efficacy and Toxicity Testing. Clin Transl Sci. 2017;10(4):237–239. doi:10.1111/cts.12444

21 Xiao S, Coppeta JR, Rogers HB, et al. A microfluidic culture model of the human reproductive tract and 28-day menstrual cycle. Nat Commun. 2017;8:14584. Published 2017 Mar 28. doi:10.1038/ncomms14584

22 Herland A, Maoz BM, Das D, et al. Quantitative prediction of human pharmacokinetic responses to drugs via fluidically coupled vascularized organ chips. Nat Biomed Eng. 2020;4(4):421–436. doi:10.1038/s41551-019-0498-9

23 Lever AR, Park H, Mulhern TJ, et al. Comprehensive evaluation of poly(I:C) induced inflammatory response in an airway epithelial model. Physiol Rep. 2015;3(4):e12334. doi:10.14814/phy2.12334

24 Huh D, Matthews BD, Mammoto A, Montoya-Zavala M, Hsin HY, Ingber DE. Reconstituting organ-level lung functions on a chip. Science. 2010;328(5986):1662–1668. doi:10.1126/science.1188302

25 Tan K, Keegan P, Rogers M, et al. A high-throughput microfluidic microphysiological system (PREDICT-96) to recapitulate hepatocyte function in dynamic, re-circulating flow conditions. Lab Chip. 2019;19(9):1556–1566. doi:10.1039/c8lc01262h

26 Azizgolshani H, Coppeta JR, Vedula EM, et al. High-Throughput Organ-on-chip Platform with Integrated Programmable Fluid Flow and Real-time Sensing for Complex Tissue Models in Drug Development Workflows. Submitted for Publication.

27 Levardon H, Yonker LM, Hurley BP, Mou H. Expansion of Airway Basal Cells and Generation of Polarized Epithelium. Bio Protoc. 2018;8(11):e2877. doi:10.21769/BioProtoc.2877

28 Mou H, Vinarsky V, Tata PR, et al. Dual SMAD Signaling Inhibition Enables Long-Term Expansion of Diverse Epithelial Basal Cells. Cell Stem Cell. 2016;19(2):217–231. doi:10.1016/j.stem.2016.05.012

29 Prachanronarong KL, Canale AS, Liu P, et al. Mutations in Influenza A Virus Neuraminidase and Hemagglutinin Confer Resistance against a Broadly Neutralizing Hemagglutinin Stem Antibody. J Virol. 2019;93(2):e01639-18. Published 2019 Jan 4. doi:10.1128/JVI.01639-18

30 Herzog, P., Drosten, C. & Müller, M.A. Plaque assay for human coronavirus NL63 using human colon carcinoma cells. Virol J 5, 138 (2008). https://doi.org/10.1186/1743-422X-5-138

31 Livak KJ, Schmittgen TD. Analysis of relative gene expression data using real-time quantitative PCR and the 2(-Delta Delta C(T)) Method. Methods. 2001;25(4):402–408. doi:10.1006/meth.2001.1262

32 Lee N, Chan PK, Hui DS, et al. Viral loads and duration of viral shedding in adult patients hospitalized with influenza. J Infect Dis. 2009;200(4):492–500. doi:10.1086/600383

33 Boivin G, Coulombe Z, Wat C. Quantification of the influenza virus load by real-time polymerase chain reaction in nasopharyngeal swabs of patients treated with oseltamivir. J Infect Dis. 2003;188(4):578–580. doi:10.1086/377046

34 Li IW, Hung IF, To KK, et al. The natural viral load profile of patients with pandemic 2009 influenza A(H1N1) and the effect of oseltamivir treatment. Chest. 2010;137(4):759–768. doi:10.1378/chest.09-3072

35 Li CC, Wang L, Eng HL, et al. Correlation of pandemic (H1N1) 2009 viral load with disease severity and prolonged viral shedding in children. Emerg Infect Dis. 2010;16(8):1265–1272. doi:10.3201/eid1608.091918

36 Hoffmann M, Kleine-Weber H, Schroeder S, et al. SARS-CoV-2 Cell Entry Depends on ACE2 and TMPRSS2 and Is Blocked by a Clinically Proven Protease Inhibitor. Cell. 2020;181(2):271-280.e8. doi:10.1016/j.cell.2020.02.052

